# Hummingbirds use wing inertial effects to improve maneuverability

**DOI:** 10.1101/2023.07.21.550104

**Authors:** Mohammad Nasirul Haque, Bo Cheng, Bret W. Tobalske, Haoxiang Luo

## Abstract

Hummingbirds outperform other birds in terms of aerial agility at low flight speeds. To reveal the key mechanisms that enable such unparalleled agility, we reconstructed body and wing motion of hummingbird escape maneuvers from high-speed videos; then, we performed computational fluid dynamics modeling and flight mechanics analysis, in which each wingbeat was resolved. We found that the birds utilized the inertia of their wings to achieve peak body rotational acceleration within half a wingbeat before the aerodynamic forces became dominant. The aerodynamic forces instead counteracted the reversed inertial forces in the other half wingbeat, thereby to sustain body rotation, albeit at a lower acceleration. Thus, individual wingbeat cycles that generated body rotations can be split into an agility phase with rapid inertial acceleration, and a stability phase with counteracting aerodynamic and inertial acceleration. This mechanism involving inertial steering enables hummingbirds to generate instantaneous body acceleration at any phase of a wingbeat, and it is likely the key to understanding the unique dexterity distinguishing them from aircraft that solely rely on aerodynamics for maneuvering.

## 1. Introduction

Hummingbirds are one of nature’s most remarkable flyers, capable of not only cruising flight like most other birds but also uniquely capable of sustained hovering and rapid, near omnidirectional aerobatic maneuvers[1]. To enable these different flight modes, their flapping wings must provide the necessary forces and moments for six degrees-of-freedom (DoFs) linear and angular accelerations, in addition to maintaining weight support. In hovering flight, hummingbirds appear to have converged evolutionarily on the wing kinematics and aerodynamics of insects[2,3]. In maneuvers, hummingbirds exhibited higher degree of aerial agility at low speeds arguably than all other birds at larger body size[4] and flying insects at similar or smaller body size[5].

Both hummingbirds and flying insects employ unsteady aerodynamics associated with their reciprocating wing movements including stroke acceleration/deceleration, wing pitching and dynamic changes of the angle of attack, and wing reversal[2,3,6,7]. In a hovering-to-escape maneuver, a Rivoli hummingbird, for example, can evade the perceived threat, reorient its body, and accelerate toward the escaping direction in less than 0.2 second or about 5 or 6 wingbeat cycles[8,9] (Fig. 1). In the process, the hummingbird body undergoes rapid pitch and roll, and is capable of reaching maximum rotational speed within a single wingbeat. How are hummingbirds able to execute such a complex maneuver (literally) in the blink of an eye? Such rapid, within-wingbeat body acceleration cannot be achieved by insects such as fruit flies which rely on subtle deviations of wing kinematics generated by steering muscle of relatively low power capacity[10– 12]. Unlike insects, hummingbirds drastically change their wing kinematics and substantially alter the aerodynamic force vector with respect to their body to generate near omnidirectional linear and rotational accelerations[8,9]. However, since their wing motion is reciprocal, the forces and torques would be reversed between half strokes and therefore could largely self-cancel. Therefore, we do not yet understand how these forces lead to the net torques needed for rapid rotational acceleration of the body. In addition, it is not clear how hummingbirds manage to control their rapid rotations without losing stability. A recent study suggests that hummingbirds use wing motion primitives that directly command the body angular rates[13], a process that promotes aerial agility and stability via simplifying flight control, similar to the “fly-by-wire” system of an aircraft[14]. However, it remains unclear how such unique traits emerge from the mechanics and neuromuscular control of hummingbird flight.

**Fig. 1.**
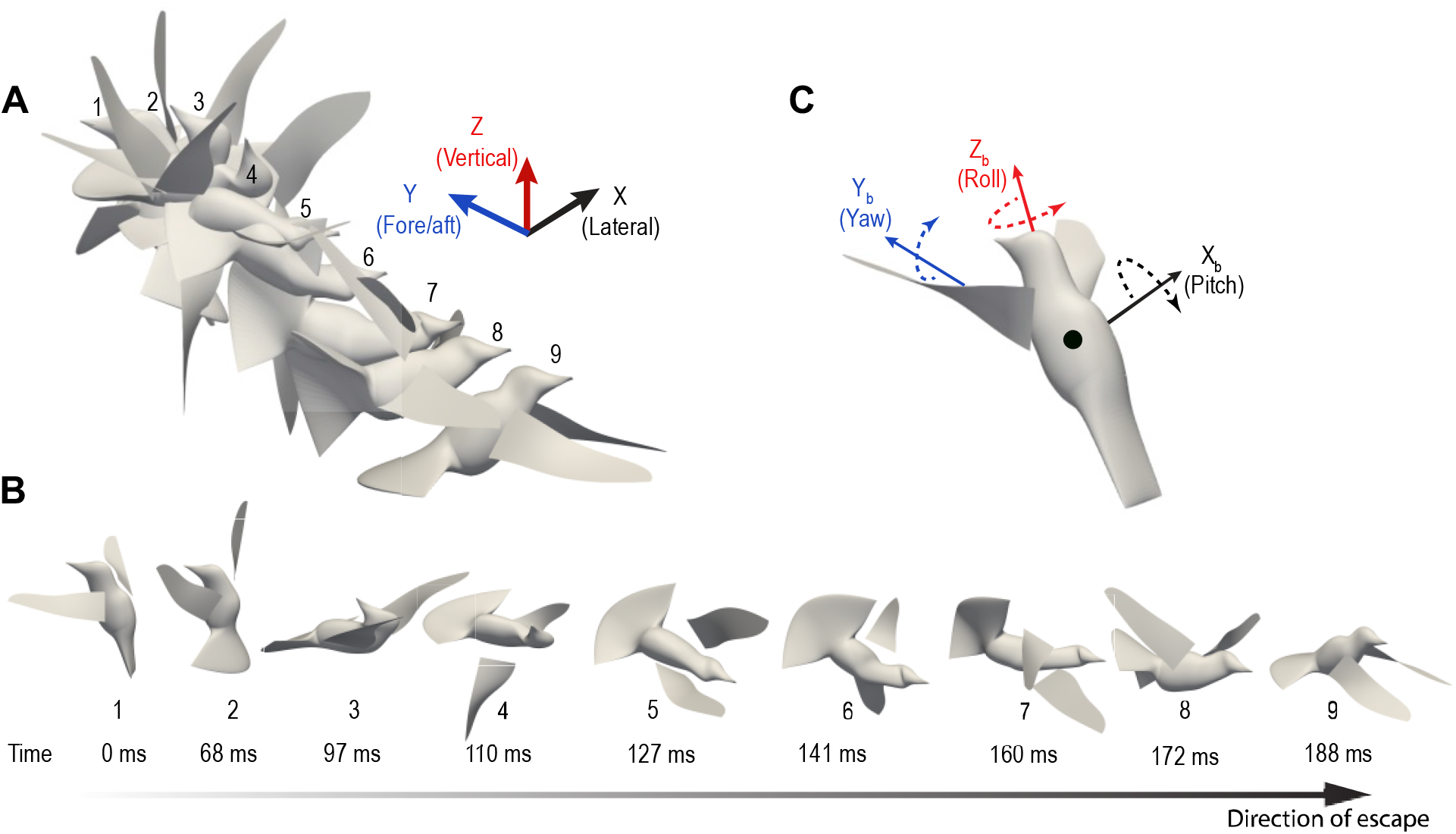
An elicited escape maneuver in a Rivoli’s hummingbird. (**A**) Snapshots of the reconstructed escape maneuver in the global reference frame; (**B**) the same snapshots separated to show body orientation; (**C**) principal axes of the bird in the body-fixed coordinate system and graphical representations of pitch, roll, and yaw rotations.

It is commonly accepted that hummingbirds, like other flying insects or man-made aircrafts, rely on aerodynamic forces to generate maneuvers. However, studies have shown that some animals are also adept at recruiting inertia of their limbs, bodies and tails for steering and stabilization, e.g., gecko[15]. Compared with flying insects, hummingbirds have relatively high wing-to-body mass ratio, and wing kinetic energy to the aerodynamic work[5], both suggesting a potentially increased role of wing inertia in maneuverability. Previous work based on wingbeat-cycle-averaged analysis typically disregard the role of inertia since the cycle-averaged inertial forces sums to approximately zero. However, the recent finding shows that hummingbirds can achieve peak acceleration as fast as in one single wingbeat[8,9], suggesting that cycle-averaged analysis may not apply to hummingbird flight and the role of wing inertia in maneuvers needs to be examined.

To address these gaps in understanding how hummingbirds accomplish their unparalleled aerial maneuverability, and to reveal the roles of aerodynamic and inertial mechanisms in generating within-wingbeat body accelerations, we analyzed the digitized high-speed videos of elicited escape maneuvers of four hummingbird species[8,9]. We first examined the timing of body pitch rotational acceleration within the wingbeat cycle to infer the primary source of torque production and form our primary hypothesis. Then, we reconstructed the full-body kinematics of the hummingbirds from the digitized pre-labelled dots and anatomical landmarks (Fig. 1), and we performed high-fidelity computational fluid dynamics (CFD) and six-DoF flight mechanics simulation of the escape process. The simulation captures both spatial and temporal details of the aerodynamic and inertial force distributions from the wings, from which we calculated the total forces and torques and revealed the mechanisms that enable the hummingbirds to perform the rapid maneuvers.

## 2. Methods

### 2.1 Experiment and reconstruction of wing kinematics

Rotational accelerations of the body were extracted from previously kinematic measures of hummingbird maneuvers in four species during elicited escape that were reported in Cheng *et al*.[8]. (Rivoli’s hummingbird, *Eugenes fulgens;* Blue-throated Mountain-Gem, *Lampornis clemenciae;* Black-chinned hummingbird, *Archilochus alexandri;* and Broad-billed hummingbird *Cynanthus latirostris*). These data were used to develop the hypotheses on inertial and aerodynamic forces that we then tested using CFD focusing primarily on flight in two male Rivoli’s hummingbirds (*Eugens fulgens*). Additional details of the original experimental design are in Cheng *et al*.[8]; briefly: Body mass (*M*) was measured using digital balance (resolution 0.01 g). The experiment was performed in an acrylic plastic transparent flight chamber with dimensions 87 × 77 × 61 cm^3^. The bird was fed via a 3-ml plastic syringe that was located at one side of the chamber. While it was hovering and feeding, the bird was startled away from the front of the feeder by a human thrusting a black clipboard toward the bird and the bird immediately flew away. The flight kinematics of the entire escape maneuver were recorded at 1000 frames/second with a shutter speed of 1/6000 sec using three high-speed video cameras, one SA3 and two PCI-1024 (resolution 1024 × 1024 pixel) (Photron Inc., San Diego, CA, USA). The bird’s body and wings were marked with 1.5 mm dots of non-toxic, water-soluble white paint prior to the experiment. A total of eighteen marker points were placed, eight points on the body and five points on each wing (Fig. S1B).

In the next step, the three-dimensional coordinates of marker points were digitized frame by frame from the videos using a custom MATLAB program[16] (Fig. S1C). To obtain more detailed kinematics for our new analysis, five additional marker points were added: one on each wing tip, one on the trailing edge of each wing, and the fifth one on the middle of the outermost tail feather. These five points were identified based on natural marks on the feathers (Fig. S1B). In the bird model for CFD, each wing was represented by a zero-thickness surface whose profile was created using spline interpolation based on the digitized marker points. The interior of the wing surface was spanned by straight chordwise segments connecting the leading edge and the trailing edge. Each wing surface consisted of 721 nodal points and 1340 triangular elements. The reconstructed wing surfaces incorporated the dynamic deformation features seen in the videos including spanwise bending and twisting. The same strategy was used previously to reconstruct hovering and cruise flight of hummingbirds[17,18].

To reconstruct the bird body, the characteristic dimensions of the bird was first extracted along the lateral and fore-aft directions within different cross sections. Based on these dimensions, a series of ellipses were used to create the cross sections from the beak to the tail, where each ellipse consisted of 80 nodes. During flight, these ellipses were transformed, including translation, rotation, and deformation, to follow the nine marker points that served as the control points (one on the bill, two eyes, one on the back, two at the body-tail junction, and three on the tail; see Fig. S1B). Such transformation incorporated changes of the body orientation as well as deformations induced by the head movement and tail flaring. The bird body including head, trunk, and tail was then meshed using the points on the ellipses. At certain stages, e.g., when the bird was turning its head to its left, the mesh around the neck was smoothed to avoid being overly distorted due to excessive strain. Overall, the body surface consisted of 15,836 triangular elements and 7,920 nodes.

About 200 frames were reconstructed (six to seven wingbeat cycles), which consisted of a short period of hovering, rapid pitch-up and roll, and linear accelerations. This stereotyped combination of pitch-roll maneuver was consistent among the bird samples across different species[8]. Thus, only two Rivoli’s hummingbirds were used for the CFD study. A temporal spline interpolation was further introduced to smooth the transition of the nodal positions in time to facilitate the small time-step simulation.

### 2.2 Morphological data and wing mass distribution in the CFD model

The morphological data of the two Rivoli’s hummingbirds were given in Table 1. The principal axes of the body were defined as in Fig. 1C along with definitions of the pitch, yaw, and roll rotations. The principal moments of inertia (MoIs) were taken from the estimates by Cheng *et al*.[9], Table A1, where the same bird samples were used for their study. The off-diagonal terms of the MoIs, i.e., 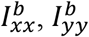, and 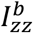were small and assumed to be zero. The MoIs were assumed to be constant during the maneuver.

**Table 1.** Morphological and kinematic data of Rivoli’s hummingbirds used in the CFD simulation.

| Parameter | Bird 1 | Bird 2 |
| --- | --- | --- |
| Mass, $M$ | 7.7 g | 7.5 g |
| Average cord length, $c$ | 1.7 cm | 1.7 cm |
| Average wing surface area, $S$ | 11.42 cm<sup>2</sup> | 11 cm<sup>2</sup> |
| Wing length, $R$ | 7.5 cm | 7.26 cm |
| Average wingtip velocity, $V$ | 11 m/s (hovering),<br>16 m/s (escape) | 11.7 m/s (hovering),<br>17.7 m/s (escape) |
| Wingbeat frequency, $f$ | 25 Hz (hovering),<br>35 Hz (escape) | 22.73 Hz (hovering),<br>37.7 Hz (escape) |
| Moment of inertia (without wings) | $I_{xx}^b = 2,119$, $I_{yy}^b = 2,741$,<br>$I_{zz}^b = 958$ g·mm<sup>2</sup> | $I_{xx}^b = 2,013$, $I_{yy}^b = 2,604$,<br>$I_{zz}^b = 910$ g·mm<sup>2</sup> |

Mass distribution for the wings was obtained by scaling the experimental data from the Ruby-throated hummingbird’s (*Archilochus colubris*, body mass *M* = 3.245 g) wings. In the experiment[19], each hummingbird wing was cut chordwise into eleven strips, and the mass of each strip was measured using a digital scale accurate to 0.0001 g, so that the one-dimensional mass distribution was obtained from the root to the tip. Since the Hummingbird’s wings consist of feathers and a musculoskeletal structure of muscle and bone, we separately distributed the mass of the bony structure along the leading edge from the root to the location of the phalanges (Fig. S2). The scaled mass data were in Table S1.

### 2.3 CFD simulation setup and verification

For the CFD simulation, the flow was considered viscous and incompressible, and was governed by the 3D Navier-Stokes equation. This equation was discretized on a single-block, nonuniform Cartesian grid and was solved with an in-house, second-order accurate immersed-boundary method[20]. A 3D domain-decomposition based parallel approach and message passing interface (MPI) were used for parallel computing. A geometric multigrid method was used for fast convergence of the Poisson equation. The computational domain was a rectangular box of size 24×21×24 cm^3^, where domain translation was adopted to follow the overall shifting position of the body (the corresponding non-inertial effect due to domain translation had been incorporated in the governing equation). For the baseline simulation, a total of 293 million (700×600×700) mesh points were used. The time-step size for the simulation was Δ*t* = 5 *μ*s. Therefore, approximately 8000 steps were used to resolve a complete wingbeat cycle during initial hovering (or about 7000 steps for later cycles). For the simulation, a total of 1000 processor cores were used on Stampede2 at the Texas Advance Computing Center (TACC).

Mesh refinement and domain size have been verified in two separate simulations to ensure that the current mesh and domain setup are adequate. To verify mesh convergence, a finer mesh was also used in simulation. Maximum grid resolution was employed around the wing during both cases. For baseline case, grid spacing was 0.03 cm in all three directions. On the other hand, grid spacing for finer grid was 0.02 cm in all three directions. Fig. S3 shows the comparison between baseline and refined mesh cases.

To verify the effect of domain size, the domain was increased in both sides of wingspan direction to repeat one wingbeat cycle simulation from t=114 ms to 146 ms for Bird 1, where the domain was increased in the Z-direction by 5 cm (i.e., box size of 24 × 21 × 34 cm^3^). Since the refined-mesh and the larger-domain simulations were done for an isolated wingbeat, rather than a continuous simulation from t=0, an addition verification of isolated cycle simulation was also performed. All these verification results were presented in Fig. S3 and showed that the baseline setup was sufficient.

### 2.4 Modeling of flight mechanics

For the simulation of the flight mechanics, the bird was considered as three separate parts, i.e., two wings and the main body, and the main body was assumed to be a rigid body going through 6-DoF movement during the maneuver. The wing-body interaction was represented by a joint force and joint torque. The linear acceleration of the main body can be written as

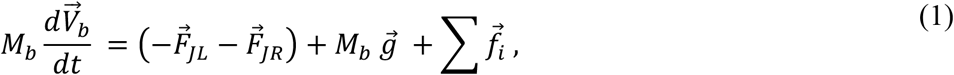

where *M*_*b*_ is the mass of the main body, 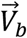 is the linear velocity, 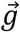 is the gravitational acceleration, 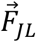 and 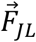 are the joint force exerted on the left and right wings, respectively, and 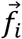 is the aerodynamic force on the body mesh nodes. The joint force was calculated by including the inertial force, gravitational force, and aerodynamic force on each wing, e.g.,

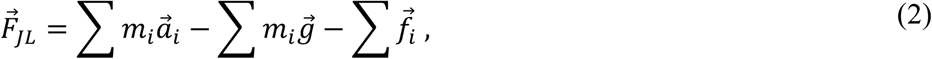

where *m*_*i*_ is mass of a mesh node on the wing, and 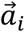 is linear acceleration of the node, and 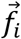 is the aerodynamic force at the node (i.e., the difference between the aerodynamic forces on the two sides of the wing).

The moment of inertia (MoI) of the main body was defined with respect to the center of mass (CoM) as ***I***_*b*_, and the rotational velocity and acceleration as 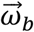 and 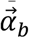, respectively. Then, the rotational dynamics of the main body can be written in the body-fixed frame as

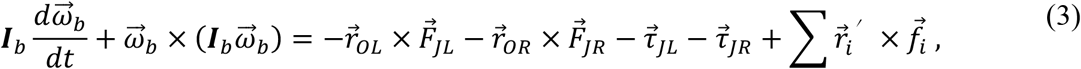

Here, 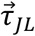 and 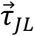 are the joint torque exerted on the left and right wings, respectively, by the main body, 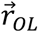 and 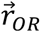 are the distance vectors from the CoM to the wing joints, and 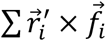 is the aerodynamic torque on the main body. The wing joint torques were calculated by including both the inertial and aerodynamic torques on the wings, e.g.,

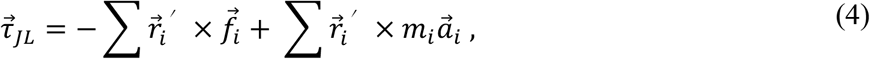

where 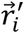 is the distance from the wing joint to the mesh node.

Once the CFD simulation was completed, the aerodynamic forces and torques on the bird’s entire body were included in the flight mechanics simulation. Eqns. (1) and (3) were integrated in time to compute the translational velocity 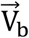 and the rotational velocity 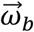.

### 2.5 Power calculation

The instantaneous aerodynamic power *P*_*A*_, inertial power *P*_*I*_, and power required for body translation *P*_*T*_, and rotation *P*_*R*_, of the bird were calculated according to the following equations. Here, *P*_*A*_ and *P*_*I*_ were calculated by integrating over the wing surfaces the product between the force at each mesh node and the velocity 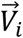 as

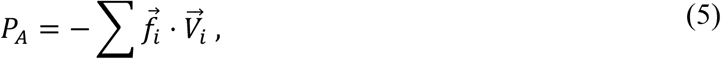

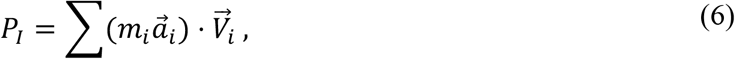

The body translational power, *P*_*T*_, and rotational power, *P*_*R*_, were calculated as follows,

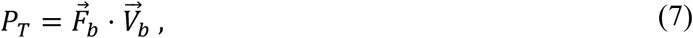

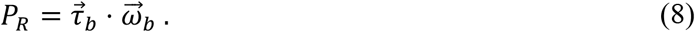

Here, 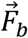 is the summation of wing joint forces from the wings to the body, and 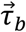 is the total torque acted on the body by the wings. These can be obtained as follows,

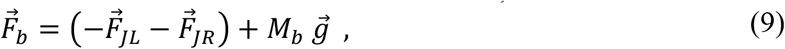

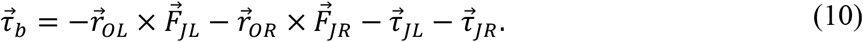

## 3. Results

### 3.1 Pitch-up rotation of four hummingbird species

When hummingbirds perceived the incoming threat in the experiment, they first performed a pitch-up rotation to lean back and then rolled the body toward either left or right side. A linear backward acceleration was also initiated. To study the mechanism of pitch rotation, we calculated the angular velocity and acceleration of the bird body in each flight trial for four species: Rivoli’s, Black-chinned, Blue-throated Mountain-Gem, and Broad-billed hummingbirds. Since the wing inertial forces and aerodynamic forces varied at different phases within one wingbeat cycle, we examined the temporal profile of the instantaneous angular pitch acceleration, *α*_*p*_ (Fig. ***2***). For all four species, the maximum angular pitch-up acceleration was nearly concurrent with pronation when the wings reached the end of upstroke and was starting the next downstroke. We termed the phase the “agility phase”, which is generally centered around pronation and has rotational acceleration. After the agility phase, pitch acceleration was significantly reduced to near zero or even reversed, although the body may continue to pitch up. The second phase generally started from mid-downstroke and may extend to part of early upstroke. We termed it the “stability phase”, as the pitch acceleration was controlled such that the pitch velocity was sustained or stabilized (i.e., not necessarily reversed with pendulum-like oscillations).

In the agility phase when body rotation accelerated, the aerodynamic forces of the wings were expected to be minimal, while the wing inertial forces were likely at their maximum (Fig. ***2***E). This observation therefore led to the hypothesis that pitch acceleration was primarily driven by the wing inertial forces within half a wingbeat, which represents an “inertial steering” mechanism. In the stability phase, the inertial forces were expected to be reversed, but the aerodynamic forces could act against the inertial forces to maintain pitch rotation initiated by the preceding inertial steering. Thus, we further hypothesized that this combined mechanism of inertial steering and aerodynamic assisting had provided the rapid and smooth body rotation during the escape maneuver.

**Fig. 2.**
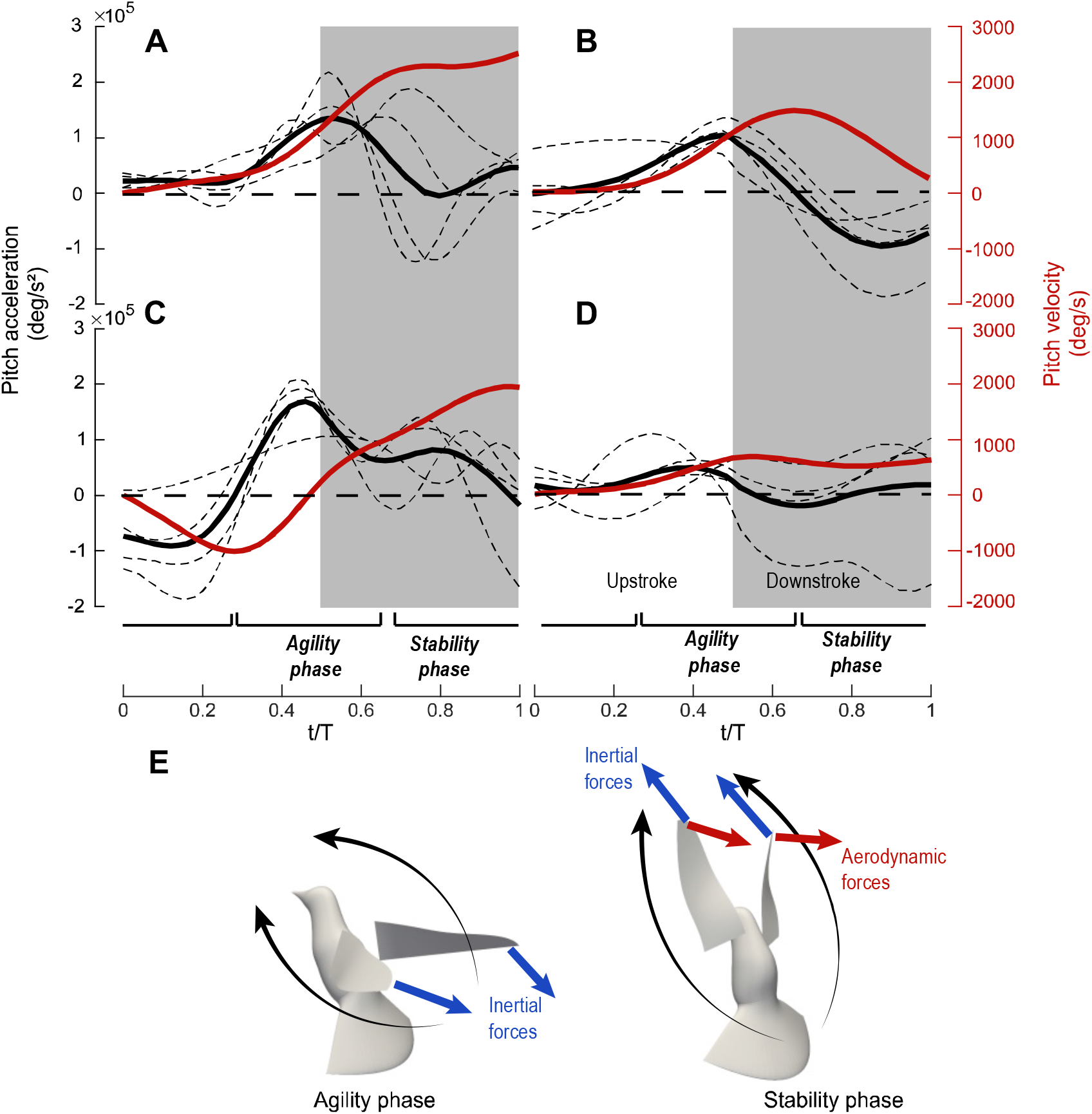
Pitch-up acceleration during the escape maneuver corresponds to the inertial torque. Subfigures show pitch acceleration values for four different species (**A**) Rivoli’s (**B**) Black-chinned Mountain-Gem, (**C**) Blue-throated, and (**D**) Broad-billed Hummingbird. Data was collected in each case from the consecutive wingbeats that correspond to the pitch-up acceleration. If multiple wingbeats were used, we phase-averaged and plotted them as a single trial case. Thin-dashed lines represent such phase-averaged acceleration of individual trials; the thick-black line represents the average among trials; the red represents the pitch velocity. For all species, maximum acceleration took place around wing pronation, which is thus termed ‘agility phase’; the rest is termed ‘stability phase’. (**E**) Illustration of aerodynamic and inertial forces on the wings at pronation and downstroke, where the forces are plotted at the wing tips for better visibility.

Species-related variations were evident in Fig. 2. Among the four species tested, Rivoli’s hummingbirds and Blue-throated Mountain-Gems underwent the most pitch, and they pitched up more than 90 degrees so that their body became upside down in the process. Black-chinned hummingbirds pitched around 75 degrees. Their pitching went through more significant oscillations within a cycle (e.g., the body acceleration reversed direction from pronation to supination), and pitching velocity was thus greatly reduced during the stability phase. As a result, this species may need three wingbeats to achieve its maximum pitching velocity, while the other three species needed only no more than two wingbeats. Broad-billed hummingbirds only pitched up by 45 degrees, and their angular acceleration was thus the least pronounced. Nevertheless, their pitch-up acceleration took place mostly during late upstroke when the aerodynamic torque likely opposed the pitching and the inertial torque most likely drove the pitching.

Guided by the hypothesis of inertial steering, we proceeded to computational modeling to examine the underlying flight mechanics.

### 3.2 Aerodynamic forces and body acceleration

The body and wing kinematics of escape maneuvers from the two Rivoli’s hummingbirds were reconstructed for CFD simulation (see Movie S1 for flow visualization). Without losing generality, the results for Bird 1 are presented here for discussion; those for Bird 2 are consistent with Bird 1 and are provided in supplementary materials.

The instantaneous aerodynamic force components changed significantly from hovering to escape (Fig. 3A; see Movie S2 for animation). During initial hovering, the stroke plane was nearly horizontal, and the wings produced weight support in both downstroke and upstroke. Around *t*=50 ms, the bird was startled and in response it flared the tail and started to escape. In the next two wingbeat cycles (*t*=60 to 115 ms), the bird tilted its stroke plane cranially relative to its body. Meanwhile, the bird body was pitching backward (pitch-up) toward the direction of escape (-Y). Due to the reorientation of the stroke plane, the aerodynamic force vector in these two wingbeats was also directed backward to initiate the linear acceleration toward the escape direction. The overall force magnitude was significantly greater than in hovering (about 1.9 times) due to a greater wingbeat frequency and thus faster stroke speed.

**Fig. 3.**
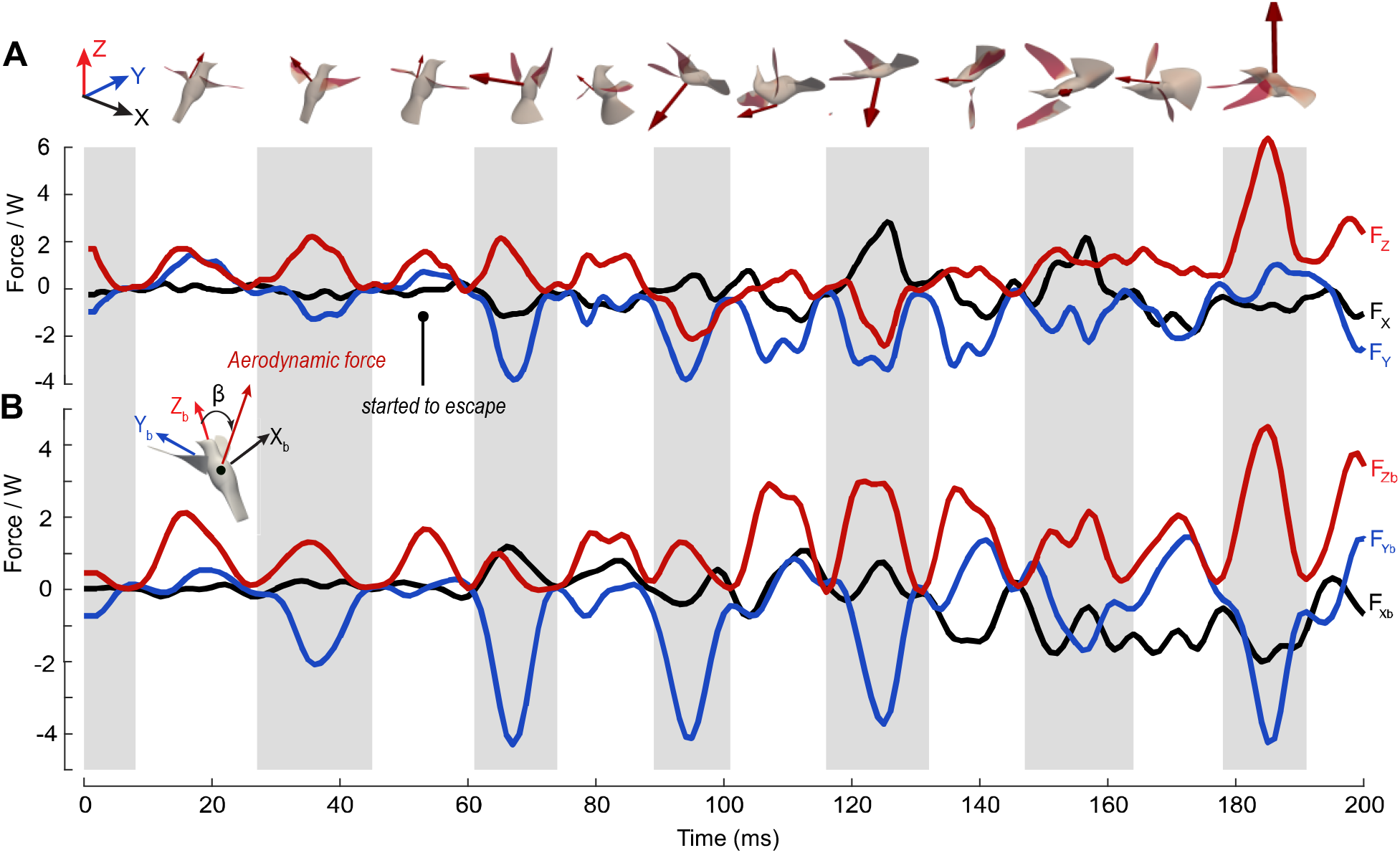
Instantaneous aerodynamic force components (normalized by body weight, *W=Mg*) on the bird (Bird 1). (**A**) In the lab frame and (**B**) in the body-fixed frame. The peak force vector is illustrated in insets for each half stroke.

In the next cycle (*t*=115 to 145 ms), the bird continued to fly in an upside-down position, while it was also rolling toward its left side. Both downstroke and upstroke produced a large force for escape acceleration. In the process, the vertical force could become negative, and consequently the bird may temporarily lose some weight support.

In the next cycle (*t*=145 to 180 ms), the bird continued to roll toward its left and was recovering from the upside-down position. The aerodynamic force vector rolled along with the body but overall continued to provide escape acceleration. Finally in the next cycle (after *t*=180 ms), the bird used a powerful downstroke to stop its body from falling and to regain its altitude. The peak force reached nearly six times of body weight.

The aerodynamic force components in the body-fixed coordinate system revealed how much the aerodynamic force vectoring changed relative to the bird body (Fig. 3B). For example, during hovering the aerodynamic force vector was around *β* = 60° from the Z_b_-axis at mid-downstroke and nearly zero at mid-upstroke; but during the first three wingbeat cycles, this angle changed to around *β* = 80° and −25°, respectively, for mid-downstroke and mid-upstroke. Such significant changes of force vectoring allowed the bird to back up away from the oncoming threat even before its body was fully rotated.

### 3.3 Inertial and aerodynamic mechanisms of pitch acceleration

After the escape maneuver was initiated, both inertial and aerodynamic torques were greatly increased compared to hovering (Fig. 4A). Overall, the inertial torque had a greater magnitude than the aerodynamic torque. In the next two wingbeats (*t* =50 to 110 ms) when pitch acceleration took place, the instantaneous net pitch acceleration was largely generated by pulsed inertial torque within the agility phase when the wings decelerated toward the end of upstroke and accelerated at the beginning of downstroke. The aerodynamic torque during the rest of the stroke cycles attenuated the opposite inertial torque that was inevitably generated by reversal acceleration of the wings, which led to the stability phase. As a result, aerodynamic torque sustained the pitch rate within a wingbeat. These results therefore supported the hypotheses we stated earlier.

**Fig. 4.**
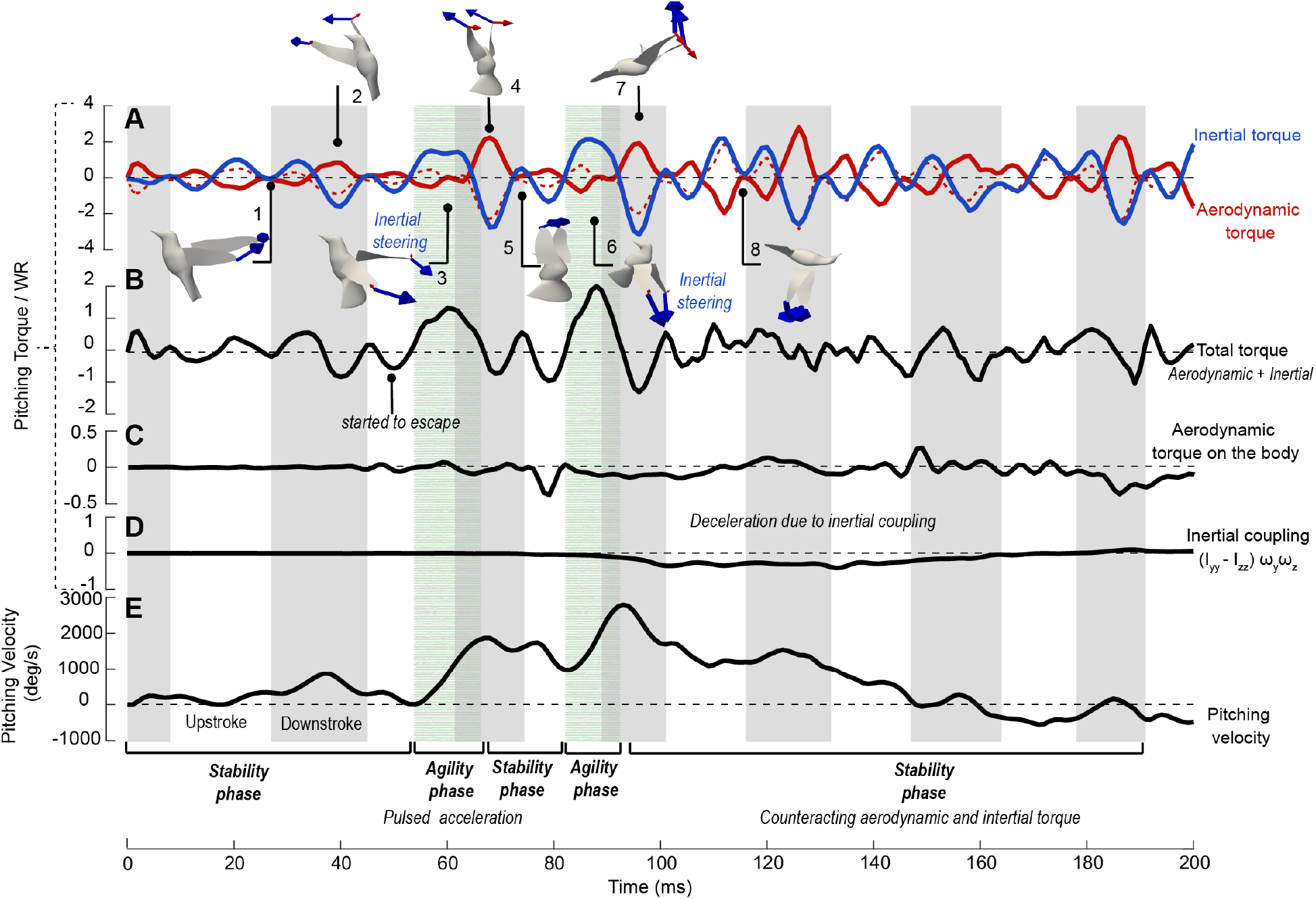
Pitching torques normalized by the body weight × wing length, *WR* (Bird 1). **(A)** The aerodynamic, inertial, and **(B)** net torques produced by the wings; **(C)** aerodynamic torque on the bird body; **(D)** the body cross-product inertial term, 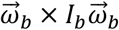; **(E)** pitching velocity of the body. Wing inertial (blue arrows) and aerodynamic (red arrows) forces were shown for selected time frames. The highlighted regions represent the agility phase where the wing inertial torque made most of contribution to the body’s pitch acceleration.

Illustrations of the wing inertial and aerodynamic forces (Fig. 4A) help us understand the torque production (see Movie S3 for a step-by-step animation in three different views). At pronation, the wings generally experienced maximum angular deceleration-and-then-acceleration. During hovering, the two wings came close to each other behind the body at pronation (Fig. 4A, inset 1). Therefore, the inertial forces of the two wings were against each other and cancelled, and thus there was no significant inertial torque at pronation. However, during pitch acceleration, the bird terminated its upstroke earlier when the two wings were still quite separated from each other at pronation (Fig. 4A, insets 3 and 6). As a result, the two inertial forces pointed backward, and they worked together to generate a torque pulse pitching up the body. After pitch acceleration, the wings resumed their stroke amplitude at pronation, and thus the inertial torque pulse disappeared for later cycles. For all the wingbeat cycles, supination did not produce a similar inertial torque pulse as pronation. This is because the two wings were close to each other at supination and their inertial forces were against each other (Fig. 4A, inset 5).

The reversed, decelerating (pitch-down) inertial torque peaked slightly after mid-downstroke (e.g., *t*=70 and 95 ms in Fig. 4A, insets 4 and 7). At mid-downstroke, the opposing aerodynamic torque was also near its maximum and thus negated this reversed inertial effect. The aerodynamic torque was highest around mid-downstroke not only because the wing velocity was the greatest at that time, but also because the stroke plane was tilted during escape and the aerodynamic force was directed backward, thus producing an increased pitch-up torque to counteract the reversed inertial torque.

The average net pitching torque of the wings for the pitch-up acceleration stage was 0.003 N-m, or 0.54 *WR*, where *W* is the body weight and *R* is the wing length, and this was significantly lower than the magnitude of individual inertial or aerodynamic peak torque (either could reach around 0.015 N-m or 2.65 *WR*). This result indicates that the pitch rotation involved intricate interplay between the instantaneous aerodynamic and inertial torques, and it was heavily influenced by the oscillatory nature of wing inertia. The wingbeat-averaged analyses on aerodynamic forces commonly used in flapping flight literature also fail to fully explain the hummingbird maneuverability[21,22].

The aerodynamic torque on the main body of the bird was overall much smaller in comparison with the wing torques (Fig. 4C). It mostly came from the flared tail, which created a resisting aerodynamic torque when the body pitched up. This torque may provide some pivoting effect at the tail so that the upper body could back away from the threat more quickly, and it may also provide dampening to help stabilize the rotation.

### 3.4 Mechanism of pitch deceleration due to inertial coupling of roll and yaw

During the pitch deceleration period (*t*=100 to 160 ms in Fig. 4A), the wing inertial forces did not produce a torque pulse. As explained earlier, the two wings resumed their stroke amplitude at pronation and their inertial effects cancelled each other (e.g., Fig. 4A, inset 8). The aerodynamic torque was mostly opposing the inertial torque, leading to a nearly zero net torque and an extended period of stability phase.

Interestingly, the cross-product term of the body rolling (*ω*_*z*_) and yawing (*ω*_*y*_) in Eqn. (3), 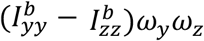, provided a significant pitching-deceleration torque (Fig. 4D). This effect was caused by the 3-DoF rotation of a rigid body, where the body yawing and rolling would contribute to the dynamics of pitching, i.e., the angular momentum of yaw was transferred to that of pitch. Magnitude of angular momentum could reach -0.26 *WR* (e.g., between *t*=100 and 150 ms). Rotational velocities also showed that there was significant overlapping between rolling and yawing for all the bird samples in terms of both magnitude and direction, consistent with this inertial coupling effect.

### 3.5 Mechanisms of roll acceleration

Both aerodynamic and inertial torques had significant contributions to roll acceleration (Fig. 5A). For both birds simulated, roll acceleration was achieved within one single wingbeat and was faster than pitch acceleration due to a lower moment of inertia for roll.

**Fig. 5.**
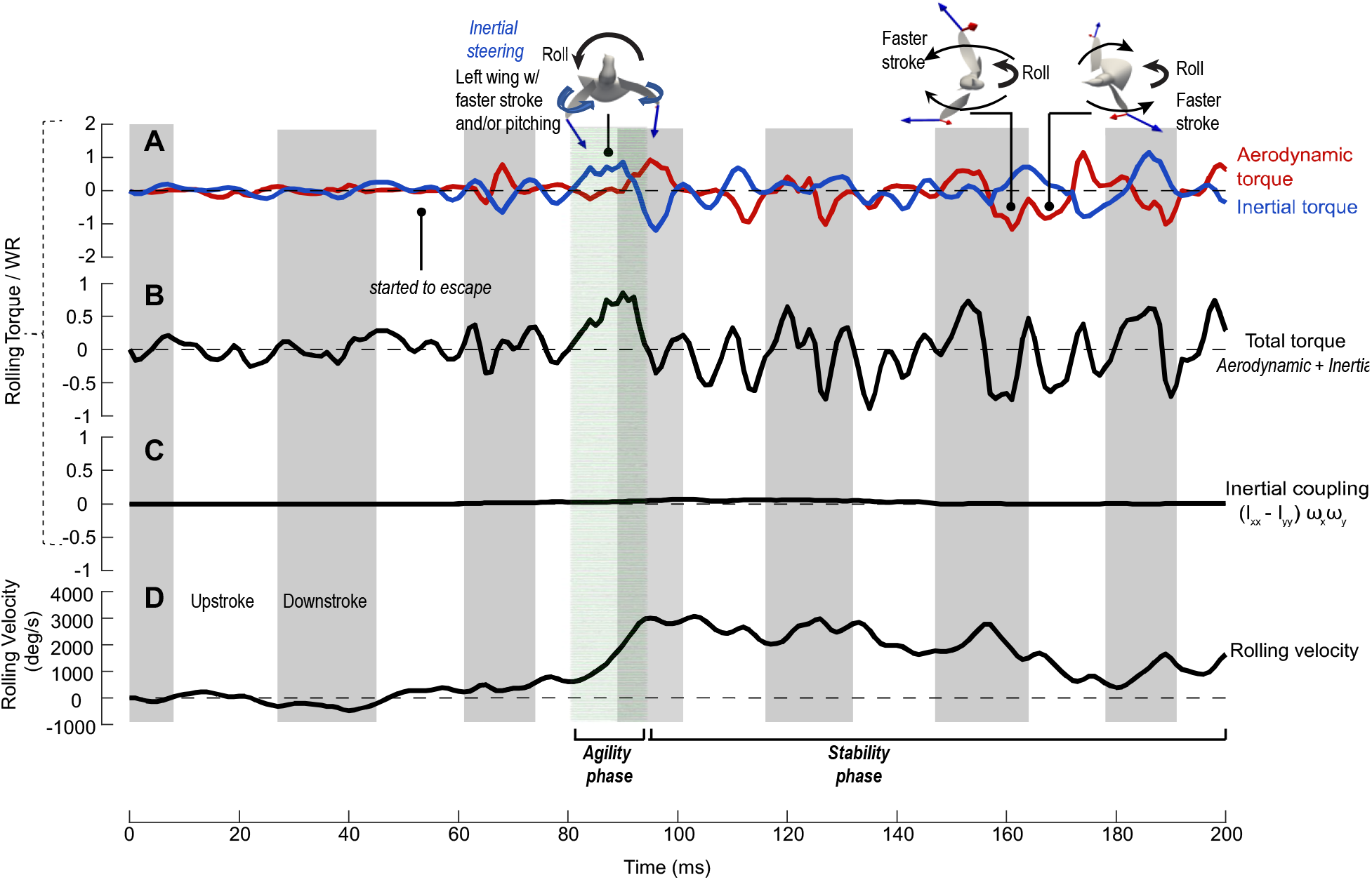
Rolling torques normalized by the body weight × wing length, *WR* (Bird 1). **(A)** The aerodynamic, inertial, and **(B)** net torques produced by the wings; **(C)** the body cross-product inertial term, 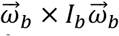; **(D)** rolling velocity of the body. Wing inertial (blue arrows) and aerodynamic (red arrows) forces were shown for selected time frames. The highlighted region represents the agility phase where the left wing had a higher momentum due to faster upstroke and/or pitching near pronation, thus creating the inertial torque for the body’s roll acceleration.

During roll acceleration (t=80 to 100 ms in Fig. 5), the net torque was mostly contributed by the positive (leftward) inertial torque during late upstroke and pronation when the aerodynamic torque was small. In the following downstroke, the large aerodynamic torque counteracted the reversed inertial torque. This inertial steering mechanism was similar to that observed in pitch.

Inspection of rolling torques on the left and right wings separately showed that the left wing had a greater stroke speed and/or a larger pitching velocity (i.e., rotation around the wing axis). As a result, the entire surface of the left wing was moving with a higher momentum than the right wing and thus produced a greater inertial force at pronation (Fig. 5, inset 1). A net inertial rolling torque was thus created. After the following mid-downstroke, the inertial torque was reversed since the left wing had to decelerate more. However, the left wing was also producing a greater aerodynamic force at this moment (Fig. S5), which led to an aerodynamic torque counteracting the reversed inertial effect. The average net rolling torque during the acceleration period, or the agility phase, was about 0.0023 N-m or 0.42 *WR*, which enabled a fast roll acceleration in about half a wingbeat.

### 3.6 Mechanism of roll deceleration due to aerodynamic damping

Once rolling was established, the aerodynamic torque became more negative (opposing roll) than positive, and the negative aerodynamic torque provided the dominant mechanism to stop the body from over-rotation during the roll-finish period (*t*=155 to 180 ms in Fig. 5). Further inspection of the velocities and aerodynamic forces of the two wings led to conclusion that such stabilizing aerodynamic torque was largely by the passive mechanism called “flapping counter-torque” [23]. Here, the flapping counter-torque was generated by combining the roll of the body and wing stroke, which led to asymmetric motion of the two wings (Fig. S5). For example, during downstroke the right wing is faster than the left wing due to the left-rolling of the body (Fig. 5, inset 2). Similarly, during upstroke the left wing is faster than the right wing (Fig. 5, inset 3). The corresponding aerodynamic drag on the two wings was also asymmetric and created a net torque against the body rolling.

### 3.7 Aerodynamic power of the maneuver

The average mass-specific aerodynamic power for the entire escape maneuvers studied here was 116.5 W/kg for Bird 1 and 124.5 W/kg for Bird 2, which was about twice of the power for hovering (46.6 W/kg and 64.3 W/kg respectively). Average mass-specific aerodynamic power for escape was 123.7 W/kg for downstroke and 105.5 W/kg for upstroke; both were approximately twice the values during hovering (64.5 W/kg and 43.5 W/kg; see Table 2 for detailed comparison). Peak aerodynamic power during escape could be substantially higher and reach more than 350 W/kg (e.g., *t* =126 and 186 ms, Fig. 6A). The combined inertial and aerodynamic power, which represents the majority of total muscle power output, may reach around 400 W/kg during downstroke for both birds.

**Table 2.** Average mass specific aerodynamic power (W/kg) during hovering and escape cycles for both birds used in the CFD simulation.

|  | Hovering power (W/kg) |  |  | Escape power (W/kg) |  |  |
| --- | --- | --- | --- | --- | --- | --- |
|  | Downstroke | Upstroke | Whole cycle | Downstroke | Upstroke | Whole cycle |
| Bird 1 | 55.7 | 38.9 | 46.6 | 126.6 | 96.8 | 116.5 |
| Bird 2 | 73.3 | 48.2 | 64.3 | 120.8 | 114.3 | 124.5 |
| Average | 64.5 | 43.5 | 55.5 | 123.7 | 105.5 | 120.5 |

**Fig. 6.**
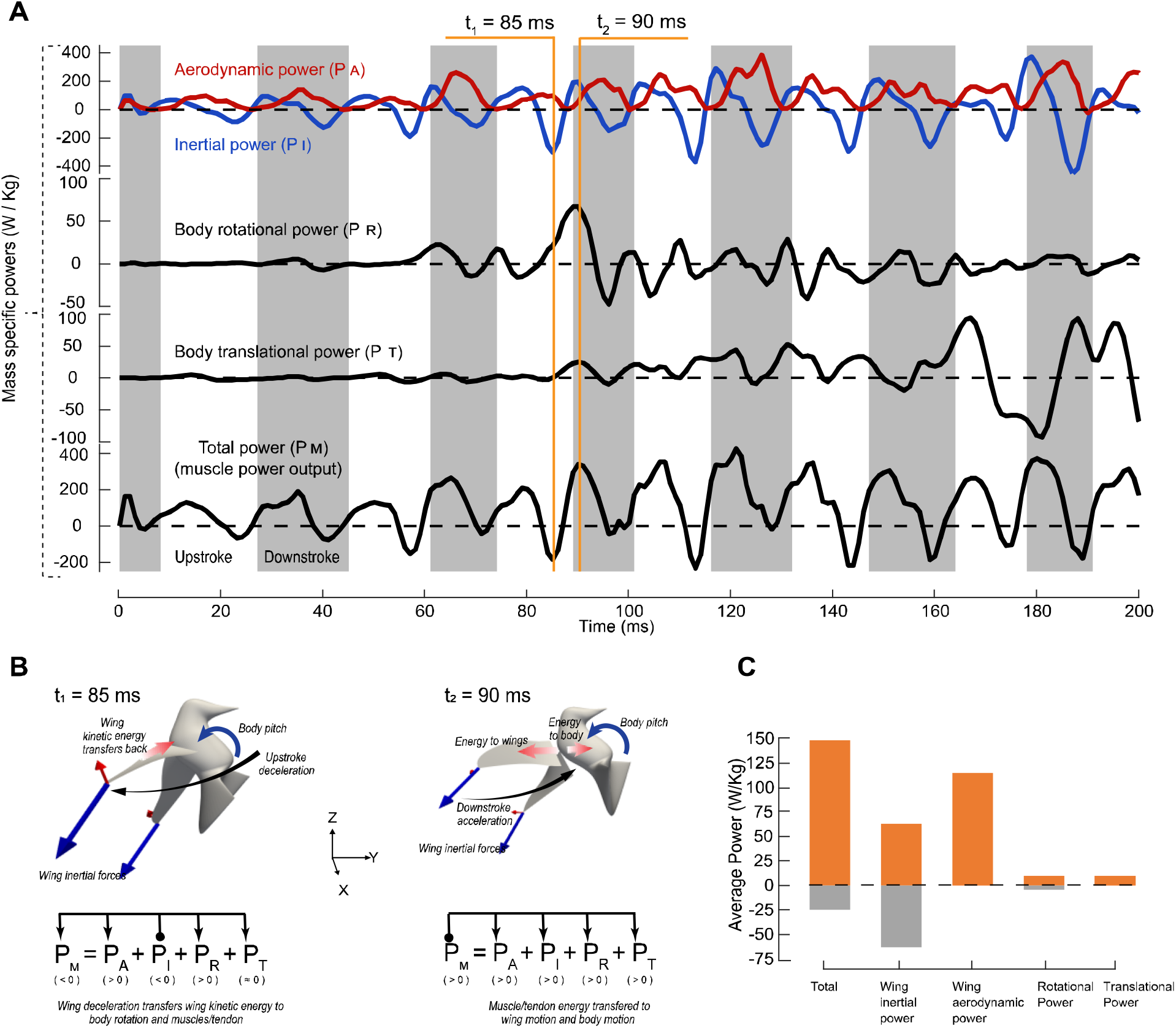
Power consumption of the escape maneuver. (**A**) Instantaneous mass-specific power of Bird 1, where the inertial power P_I_ and aerodynamic power P_A_ of the wings, the body rotational power P_R_ and translational power P_T_, and total muscle power output P_M_ were plotted. (**B**) Illustration of power flow during inertial steering. (**C**) Average positive and negative power in a wingbeat cycle.

The total power output from the muscles at any time moment, P_M_, was split among the wing inertial power, P_I_, the wing aerodynamic power, P_A_, power for the body’s rotation, P_R_, and power for the body’s translation, P_T_ (Fig. 6A), so that P_M_ = P_I_ + P_A_ + P_R_ + P_T_. For flapping wings in general, the wing inertial power is positive when the wings are accelerating and thus gaining kinetic energy; conversely, the wing inertial power is negative when the wings are decelerating and releasing the kinetic energy. All or part of this negative inertial power is converted to the aerodynamic power if the wings move against significant aerodynamic forces at the time; thus, the muscle outpower is reduced. The remaining negative power, if any, could potentially be stored in the wing muscle system[24,25]. Overall, the wing inertial power sums up to zero over a wingbeat.

During escape, the magnitude of the wing inertial power was significantly increased as compared with that during hovering, and this included both positive and negative inertial power (Fig. 6A). However, during downstroke most of the negative wing inertial power was transferred to the aerodynamic power, and the muscle output power was mostly positive. This feature may be beneficial since the negative energy may not be all stored by the muscular system[26].

Significant negative total power still existed toward the end of upstroke since the aerodynamic power was not high enough to absorb the negative wing inertial power. Yet, by transferring the majority of the negative inertial power to aerodynamic power, the cycle-averaged positive total wing actuation power is only moderately higher than the aerodynamic power (Fig. 6C). In comparison with the wing inertial or aerodynamic power, the body rotation and translation powers were much lower (Fig. 6A and C). Furthermore, the body rotation only required power input temporarily during the rotational acceleration. The sources of rotational power could come either from the negative inertial power, or from the muscle output, as explained below.

For example, at t_1_ marked out in Fig. 6A (t=85 ms) when inertial steering was happening, the wings were decelerating toward the end of upstroke (Fig. 6B), and the body was pitched up due to the wing inertial forces as discussed earlier. As a result, the negative wing inertial power due to wing deceleration (P_I_<0) was partially transferred to pitch rotation of the body in addition to transferring back to muscles (i.e., P_R_>0 and P_M_<0). At t_2_ (t=90 ms) when the wings were starting downstroke, the inertial power was positive (P_I_>0) due to wing acceleration. The reaction forces at the shoulder joints continued to pitch up the body. Thus, the muscle power at this time was output to simultaneously power wing acceleration and body pitching (Fig. 6B) with P_M_, P_I_, P_R_ all being positive. These results indicate that the bird’s inertial steering, which was enabled by the angular momentum transfer from the wings to the body as discussed in the torque analyses, was also partially supported by the energy transfer from the wings to the body.

## 4. Further Discussion

As pointed out by Cheng *et al*.[8], hummingbirds change wing kinematics significantly relative to their body from hovering to the escape maneuver, which leads to corresponding large alterations of aerodynamic force vectoring with respect to their body axes. In addition to confirming this result through computational fluid dynamics simulation of the detailed aerodynamics, the present study has provided novel insight about how the hummingbirds utilized a combination of the inertial and aerodynamic forces associated with reciprocating wing strokes to generate the force moments for fast and well-controlled body rotation. Contrary to the perhaps intuitive thought that the aerodynamic forces create the torques needed to drive the pitch, roll, and yaw rotation of the body during a maneuvering flight, our results revealed that the torques for driving the pitch and roll acceleration in an escape maneuver were more from the wings’ inertia forces than from the wings’ aerodynamic forces; the aerodynamic torques provided the counteracting effect against the opposite inertial torques due to reversal of the wing stroke. Consequently, the wing stroke responsible for pitch and roll acceleration could be roughly divided into two different phases. One was an “agility phase”, which was around the pronation and included late upstroke and early downstroke. In this phase, the inertial torques from the wings drove the body’s rotational acceleration. The second was a “stability phase”, which included mid-downstroke to early upstroke. In this phase, the aerodynamic torques counteracted the reversed wing inertial torques and helped maintain the pitch and roll velocities. With these two phases, the hummingbirds achieved great agility for rapid rotations.

The high magnitude of the wing inertial forces provided short burst moments needed for fast body rotational acceleration. For example, the inertial rolling torque from the wings was around 0.42 *WR* during roll acceleration. Given that the roll moment of inertia of the body was about 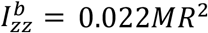, the burst torque led to a roll acceleration of more than 2000 deg/s within 15 ms. For pitch acceleration by wing pronation, the inertial torque was around 0.88 *WR*, which led to an acceleration of near 2000 deg/s within 13 ms for the pitch moment of inertia at 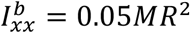. The wing inertia thus provided a great “inertial steering” mechanism for the hummingbirds to achieve rotational agility.

Inertial steering is also used by some other animals for maneuvering and stability in aerial actions. For example, a falling gecko relies on its tail to perform rapid, zero-angular momentum air-righting and turn the body[15]. The hawkmoth moves its abdomen dynamically to pitch the thorax and thus redirects lift forces for effective flight control[27]. The rose-breasted cockatoo changes the moment of inertia of its wings through wing flexing to generate net inertial torque in a stroke cycle for a roll reorientation in a flapping, banked turn[28]. However, unlike those animals that use the inertia of other body parts like tail or abdomen, the hummingbirds produced inertial steering through their wings while they were in flapping motion. In addition, unlike cockatoos, the hummingbirds did not change the wings’ moment of inertia to generate the net inertial torque, which would require drastic changes of the wing shape. Instead, the hummingbirds took advantage of the favorable inertial torque within one wingbeat and cancelled out the unfavorable inertial torque by synchronizing it with the favorable aerodynamic torque.

Flying insects, as well as aircraft, utilize aerodynamic mechanisms for flight maneuvers. For example, fruit flies employ subtle deviations from hovering wing kinematics[10–12] to alter lift and drag and generate steering force moments for body rotations in saccade. We interpret that this is due, at least in part, to the low inertia of their body and limitation of their sensorimotor system, as accelerations generated by large deviations would destabilize their flight and their sensorimotor system may not respond rapidly enough[29]. Fixed-wing airplanes use elevators, rudders, and ailerons to generate aerodynamic moments for pitch, yaw, and roll controls. Helicopters and multirotor drones maneuver by adjusting the aerodynamic lift of rotors. The novel use of inertial steering for flight maneuvers by the hummingbirds should therefore be an inspiration for those seeking to develop highly maneuverable aerial vehicles.

