## Supplemental figures for "Hummingbirds use wing inertial effects to improve maneuverability"

—

**This PDF file includes:**

Supplementary Text
Figs. S1 to S10
Tables S1
Captions for Movies S1 to S3

**Other Supplementary Materials for this manuscript include the following:**

Movies S1 to S3

### Model validation

The simulated translational velocities and rotational velocities based on the CFD results were compared with those velocities derived from the digitized marker point positions. The comparison for Bird 1 was shown in **Error! Reference source not found.**A for translation and **Error! Reference source not found.**for rotation. The major velocity components for translation were the escape velocity in the Y-direction (i.e., fore-aft velocity) and the vertical velocity; both generally agreed with the experiment data. From the time around $t$=50 ms when the bird responded to the stimuli, to around $t$=160 ms when the bird finished its body rotation, the bird's fore-aft velocity went from zero to -2 m/s with an average linear acceleration close to 1.9G. The vertical velocity was mostly negative and reached around -1 m/s, indicating some loss of weight support during the maneuver. In comparison, the lateral drift velocity was much smaller. Bird 2 went through a similar fore-aft acceleration but had less vertical dropping.

Fig. S4B shows that the simulation also captured the overall characteristics of the pitch and roll rotations during the maneuver. Specifically, the pitch-up motion between $t$=50 to 170 ms, and the left-roll motion between $t$=80 and 170 ms, were evidently captured by the simulation. There were differences in the comparison, which were most likely due to the limitations of our model, e.g., the assumption of the mass distribution of the wings and body, as well as the rigid-body assumption for the head-trunk combination. In terms of magnitude, the pitching and rolling velocities may reach around 3000 deg/s, and yawing around 1500 deg/s in the middle of escape. There was significant time-overlapping among these three rotational velocities. Bird 2 had relatively lower rotational velocities (around 2000 deg/s for pitch and roll and 1000 deg/s for yaw). In terms of the sequence of rotations, Bird 2 started left rolling almost immediately after pitching up. Unlike Bird 1 who was flying on its back during maneuver, Bird 2 was flying on its left side for the next 3 wingbeat cycles after pitch-up.

### Mechanisms of yaw rotation

The overall magnitudes of the wing inertial and aerodynamic yawing torques were comparable to each other (Fig. S11), and they were close to those of the rolling torques. In addition to the wing inertial and aerodynamic effects, the cross-product body inertial term, was around -0.17$WR$ on average, which made significant contribution for yaw acceleration. This result suggested that the yaw could be caused by combination of the pitch and roll rotations, i.e., the angular momentum being transferred from pitch to yaw. To control the yaw rotation (t=145 to 180 ms), the net wing torque (around 0.49 $WR$) was contributed mostly by the aerodynamic torque that stopped the body from further yawing.

### Summary of the CFD results for Bird 2

Bird 2 had similar force characteristics as Bird 1 (Fig. S7). That is, not only the force magnitude was much greater than hovering, but also the force vector changed significantly in the body-fixed coordinate system and was mostly directed to the escape direction. However, this bird was not completely upside down and did not lose as much weight support in the process.

The results of rotational dynamics for Bird 2 were provided in Fig. S8 to S9. The general characteristics of the torque components were similar for the two birds, although there were some noticeable variations. From these results, For Bird 2 the inertial torque contribution to the pitch acceleration was even more pronounced. On the other hand, the cross-product term still helped with pitch deceleration even though its effect was not as strong as in Bird 1.

For the rolling motion of Bird 2, both aerodynamic and inertial torques contributed significantly to roll initiation, while the inertial torque was dominant in roll initiation for Bird 1. Further inspection of the left and right wing torques (Fig. S9) showed that the aerodynamic force (mostly lift) of the left wing at upstroke helped provide a left-rolling torque for roll acceleration. Another difference was that for Bird 2, yaw acceleration was contributed by both the inertial torque and the body's inertial coupling term, while for Bird 1, yaw acceleration was mostly due to the inertial coupling.

**Fig. S1.**

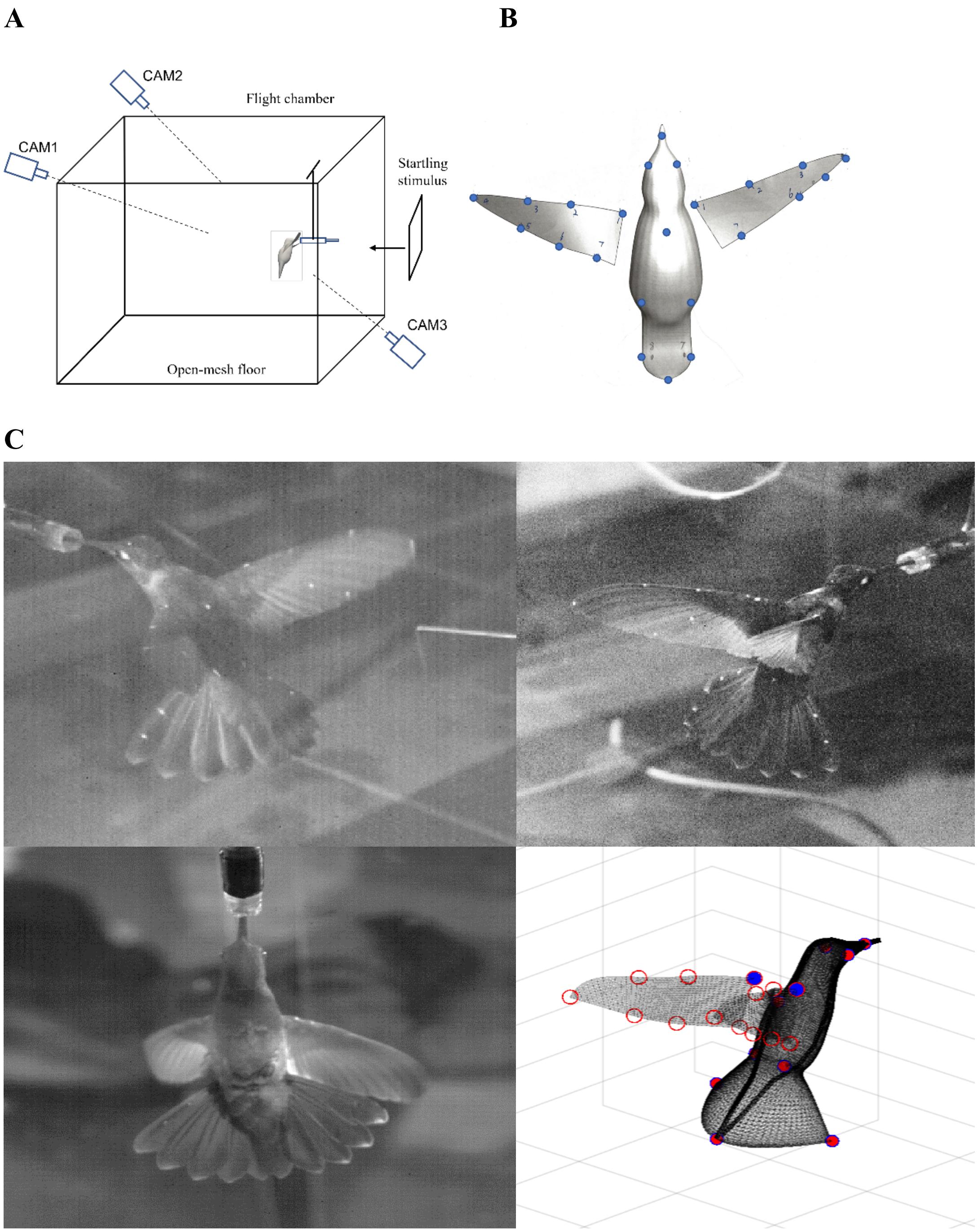


**Experimental set-up and reconstruction of the bird's model from high-speed video. (A)** Schematic of flight chamber and approximate position of three cameras used in the experiment. (**B)** Illustration of the marker points on the head, body, wings, and tail of the bird. (**C)** Snapshots from the three camera views and corresponding reconstructed bird model.

**Fig. S2.**

**A**

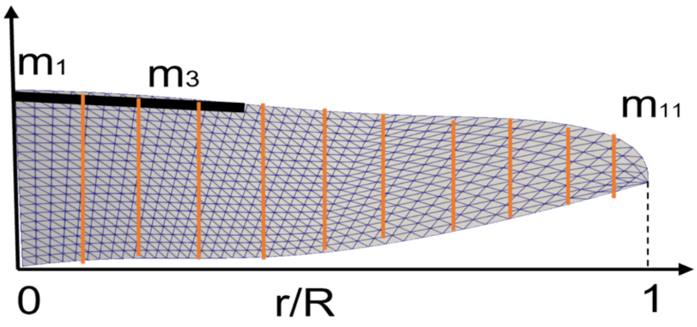


**B**

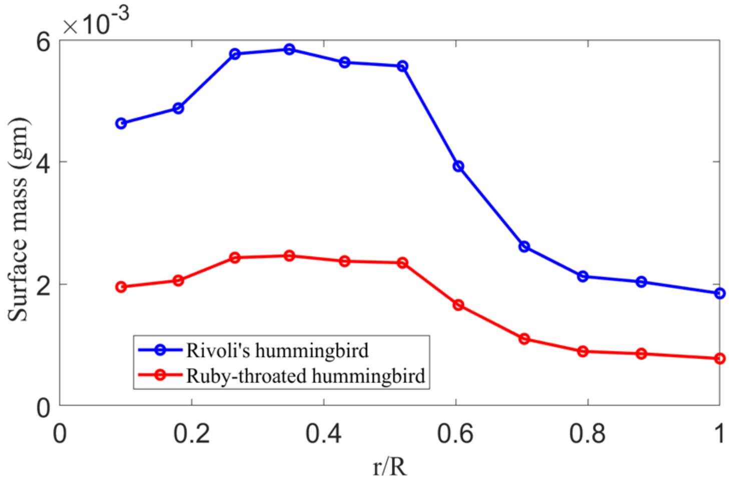


**Mass distribution of the Rivoli's hummingbird wing scaled from available data for the Ruby-throated hummingbird.** (**A**) Wing segments measured for mass. (**B**) Wing surface mass of each segment.

**Table S1. Wing mass of Rivoli's hummingbirds scaled from the Ruby-throated hummingbird.**

| **Parameter** | **Rivoli's Hummingbird 1** | **Rivoli's Hummingbird 2** | **Ruby-throated hummingbird** |
|---|---|---|---|
| Total Mass | 7.7 g | 7.5 g | 3.245 g |
| Single wing mass | 0.1962 g (scaled) | 0.1911 g (scaled) | 0.0827 g |

**Fig. S3.**

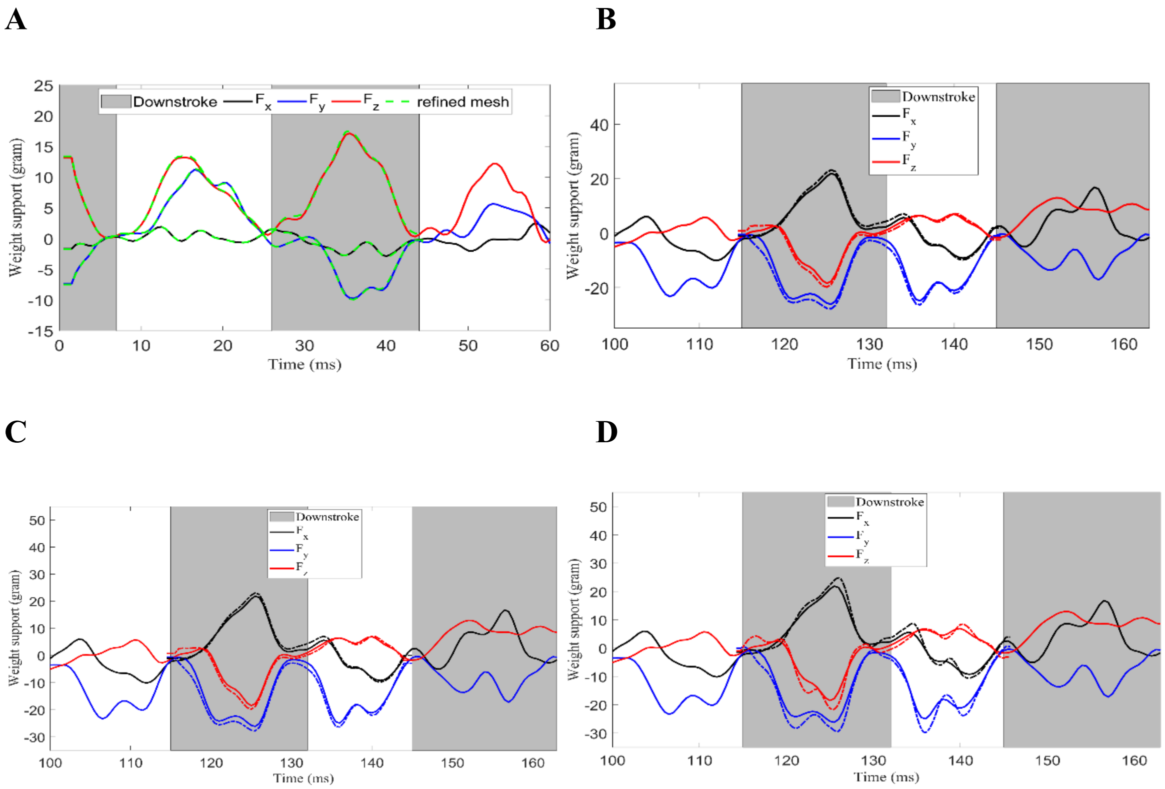


**Verification of the CFD model.** (**A**) Comparison of aerodynamic forces between the baseline and refined-mesh simulations for hovering (t=0 to 45 ms). (**B)** Comparison of aerodynamic forces between the baseline simulation and an isolated wingbeat cycle (dashed line) from t=115 to 145 ms. (**C**) Comparison of aerodynamic forces between the baseline simulation and the isolated cycle (dashed line) with a larger domain. (**D)** Comparison of aerodynamic forces between the baseline simulation and the isolated cycle (dashed line) during escape with refined mesh.

**Fig. S4.**

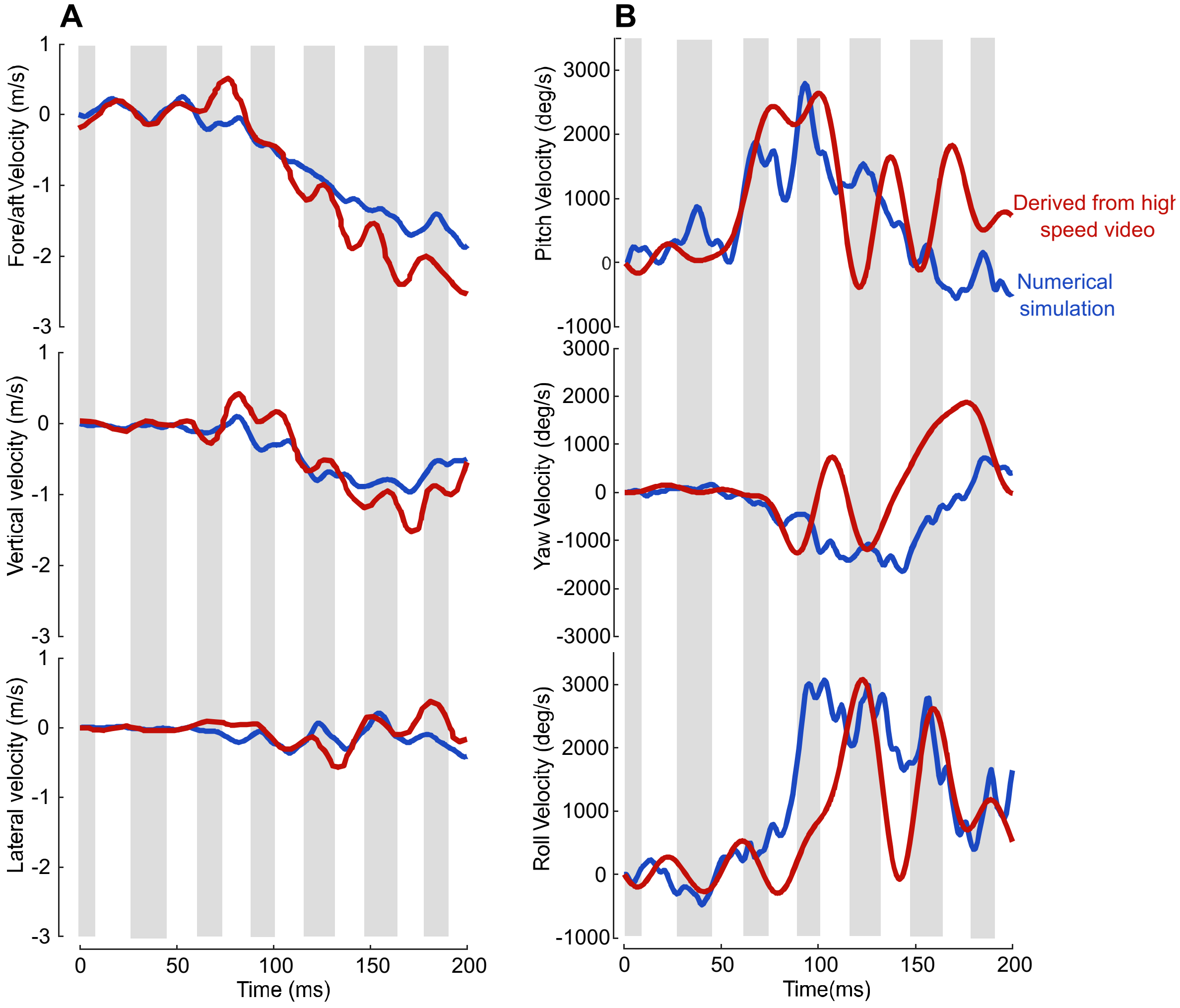


**Validation of the CFD model.** Comparison of the body's (**A**) translational velocities and (**B**) rotational velocities of Bird 1 between the CFD-based simulation and the experiment results.

**Fig. S5.**

**A**

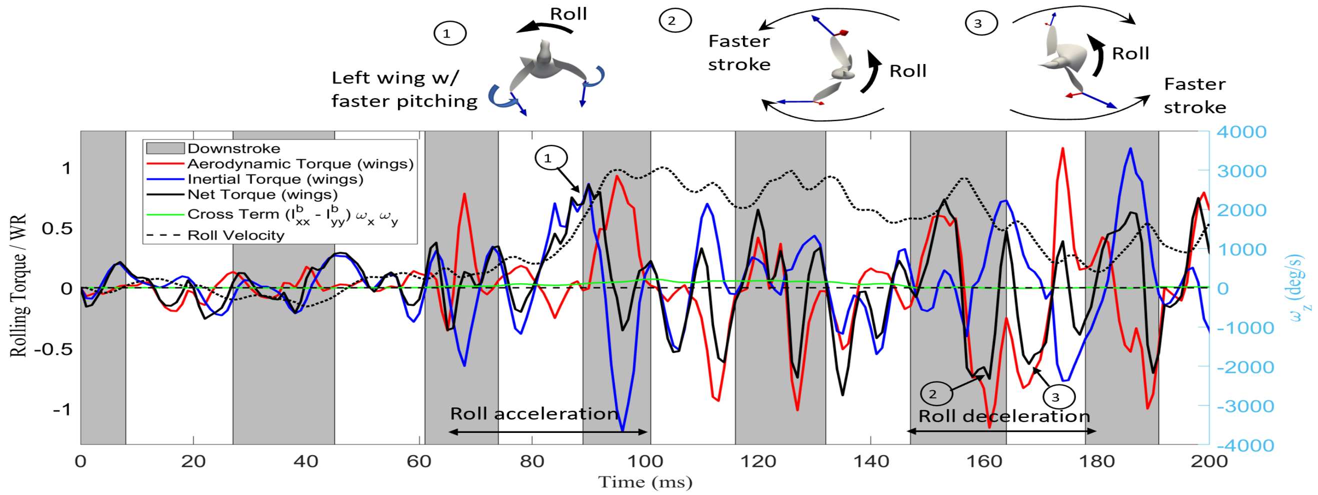


**B**

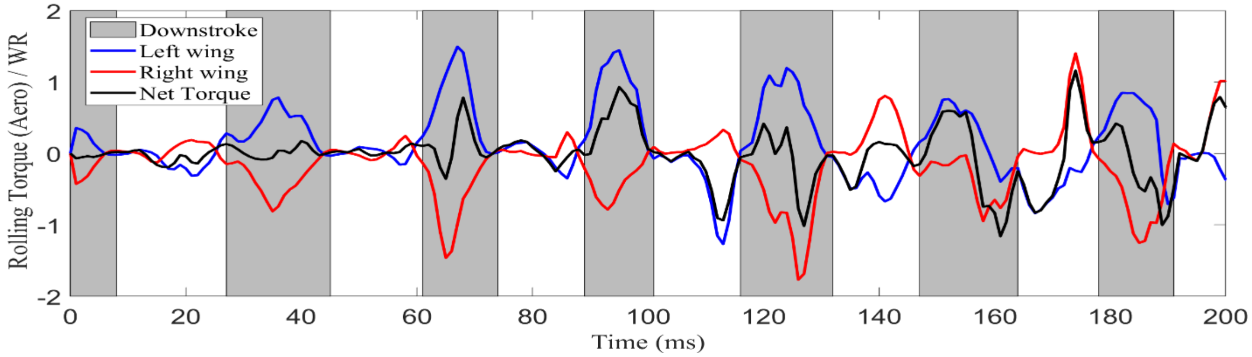


**C**

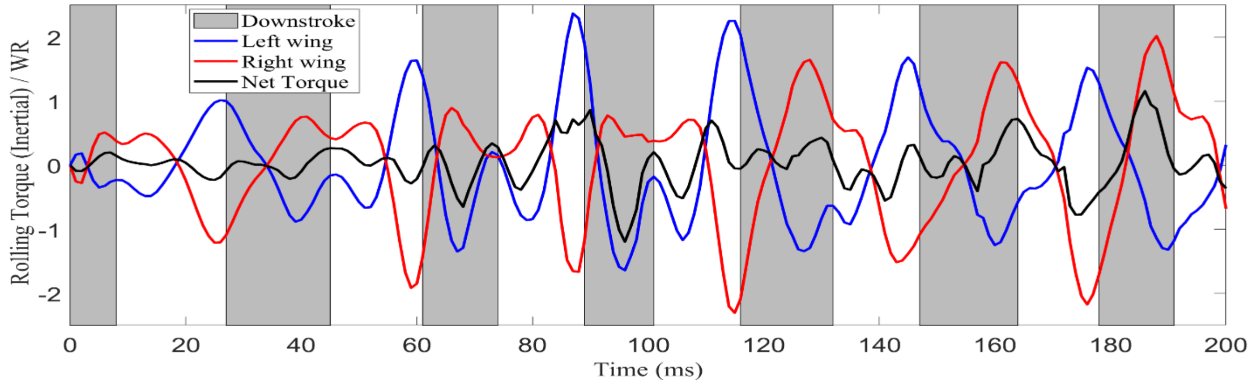


**Rolling torques (normalized by *WR*) for Bird 1**. (**A**) Rolling torques generated by both wings. (**B**) Aerodynamic rolling torques produced by left and right wings**,** (**C**) Inertial rolling torques produced by left and right wings.

Fig. S6.

A

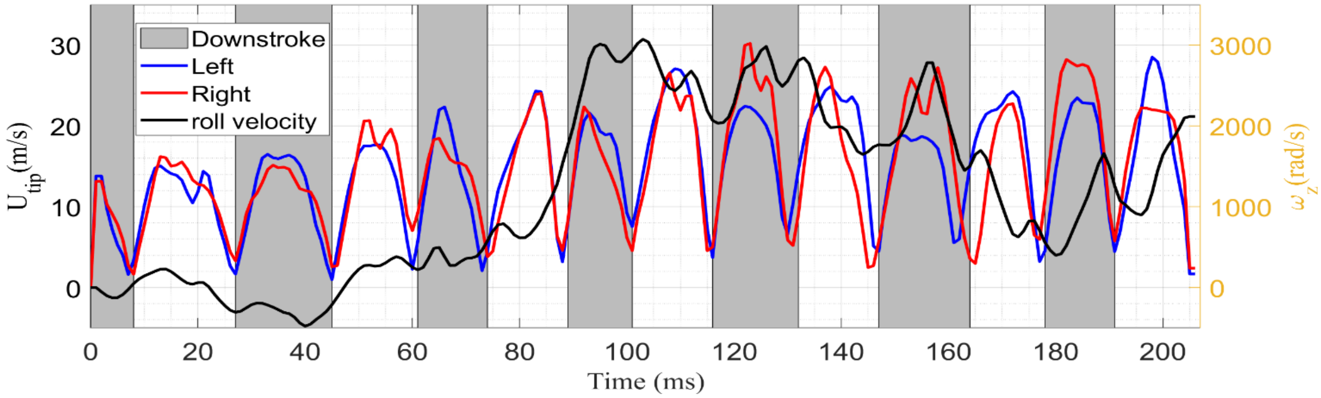


B

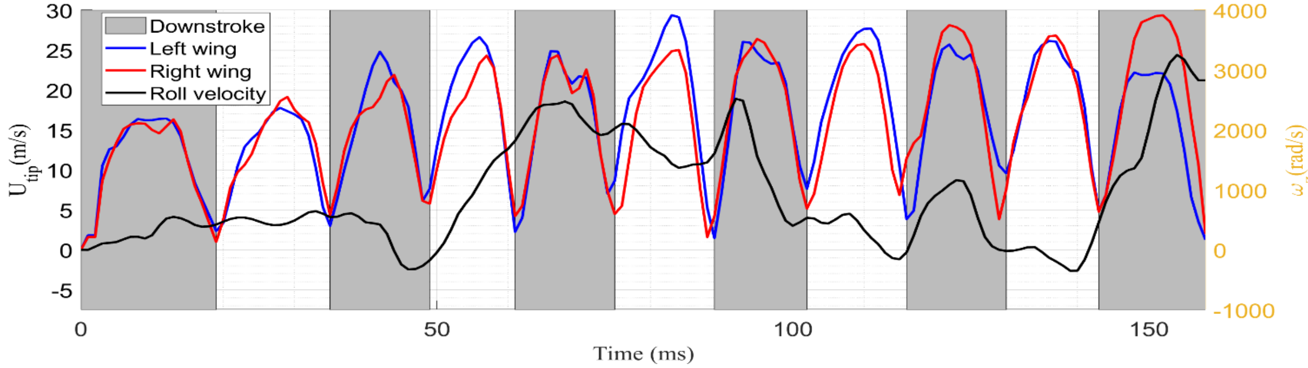


**Instantaneous wingtip velocity, $U_{tip}$, from left and right wings and roll velocity, $\omega_z$, of the body.** (**A**) Bird 1**,** (**B**) Bird 2.

**Fig. S7.**

**A**

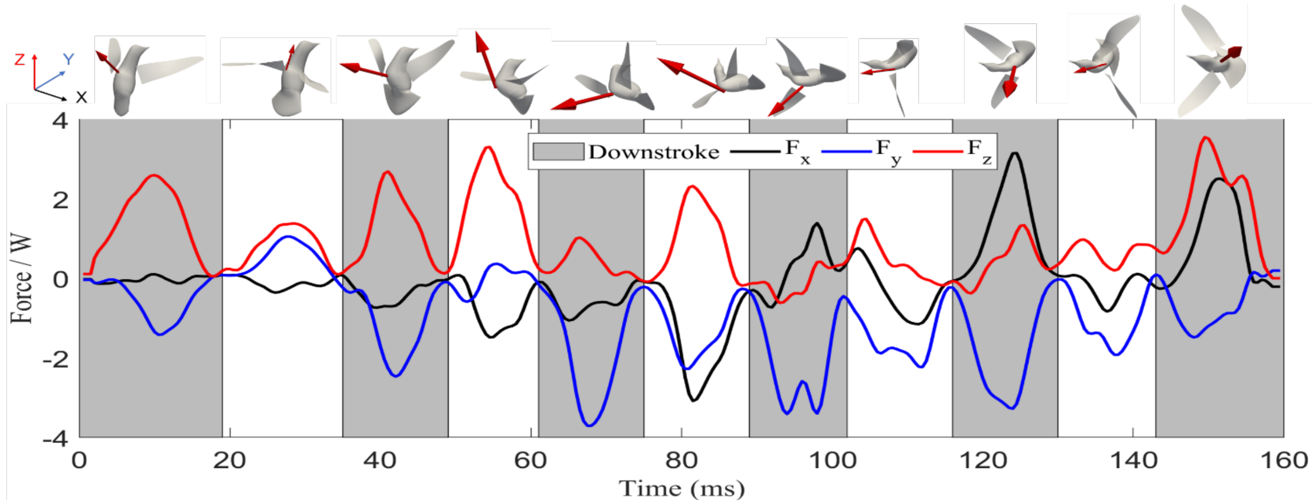


**B**

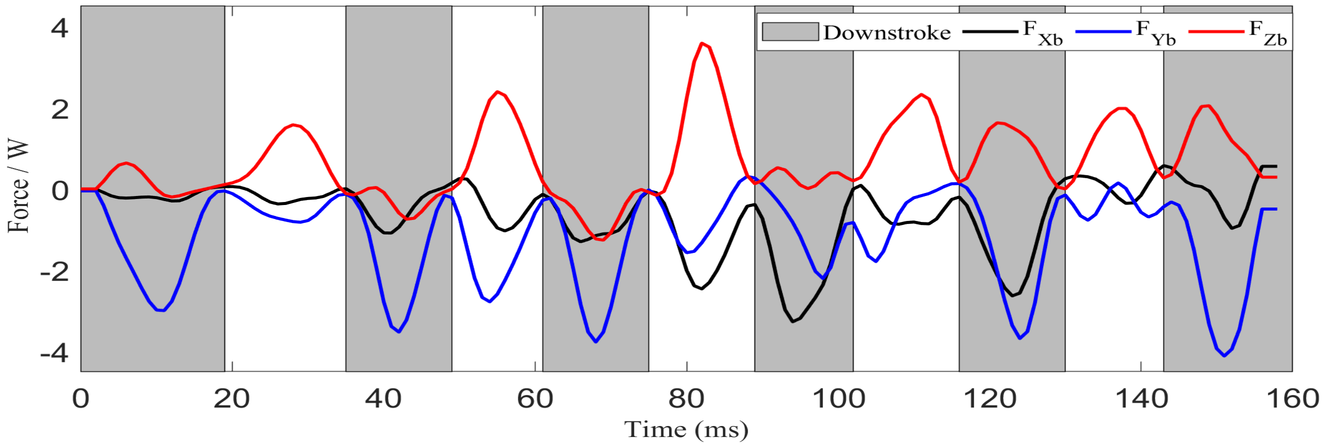


**Instantaneous aerodynamic force components (normalized by body weight,** W) **for Bird 2.** (**A**) Force components in the global frame. (**B**) Force components in the body frame. The peak force vector was illustrated for each half stroke.

**Fig. S8.**

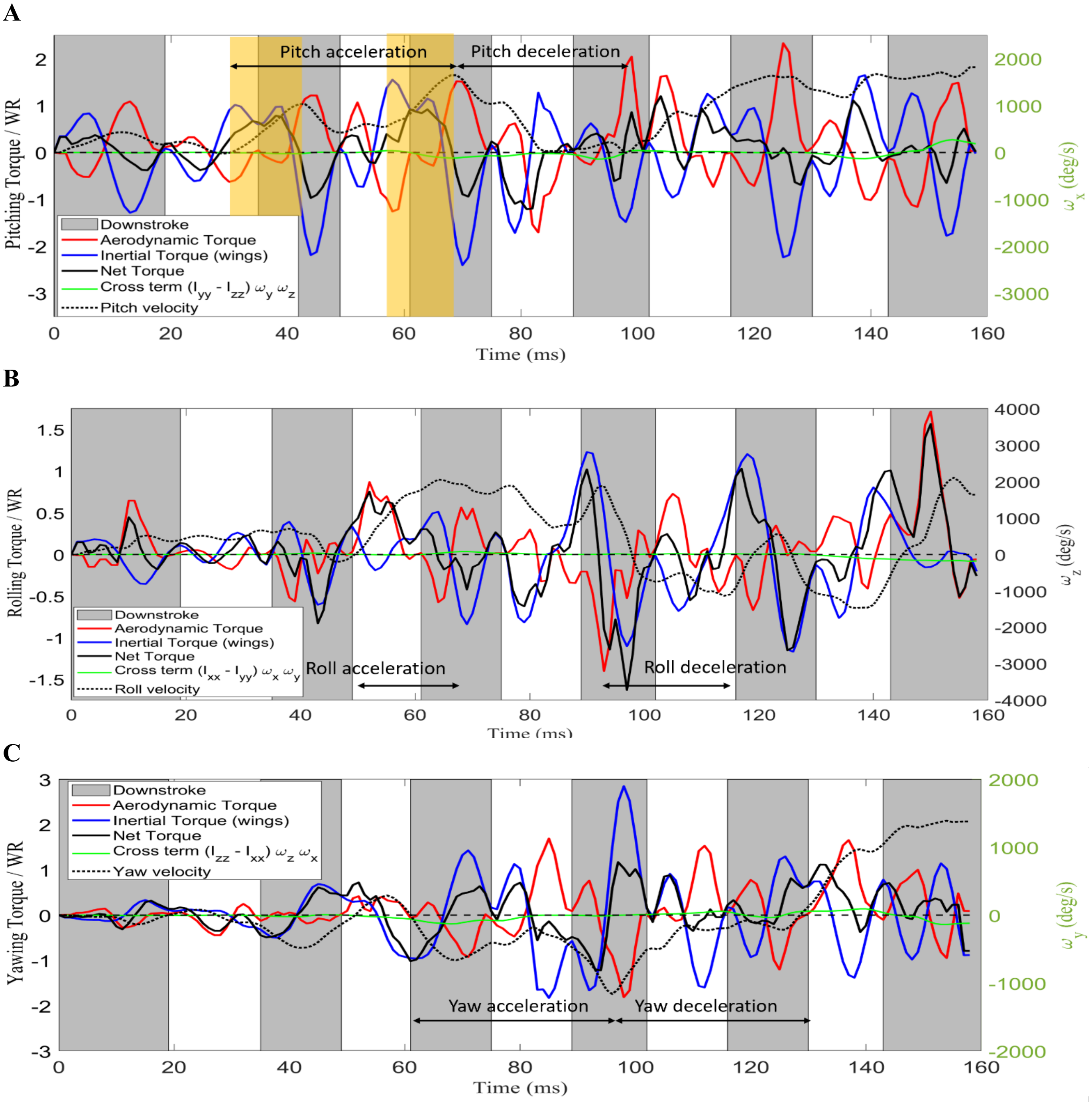


**Torques on the body for Bird 2.** (**A**) Pitching, (**B**) rolling, (**C**) yawing torques (normalized by *WR*), where the aerodynamic, inertial, and net torques produced by the wings were plotted. The body cross-product inertial term, $\vec{\boldsymbol{\omega}}_{\boldsymbol{b}} \times \boldsymbol{I_b}\vec{\boldsymbol{\omega}}_{\boldsymbol{b}}$ was also included. In **A**, the highlight regions represent inertial steering for pitch-up.

**Fig. S9.**

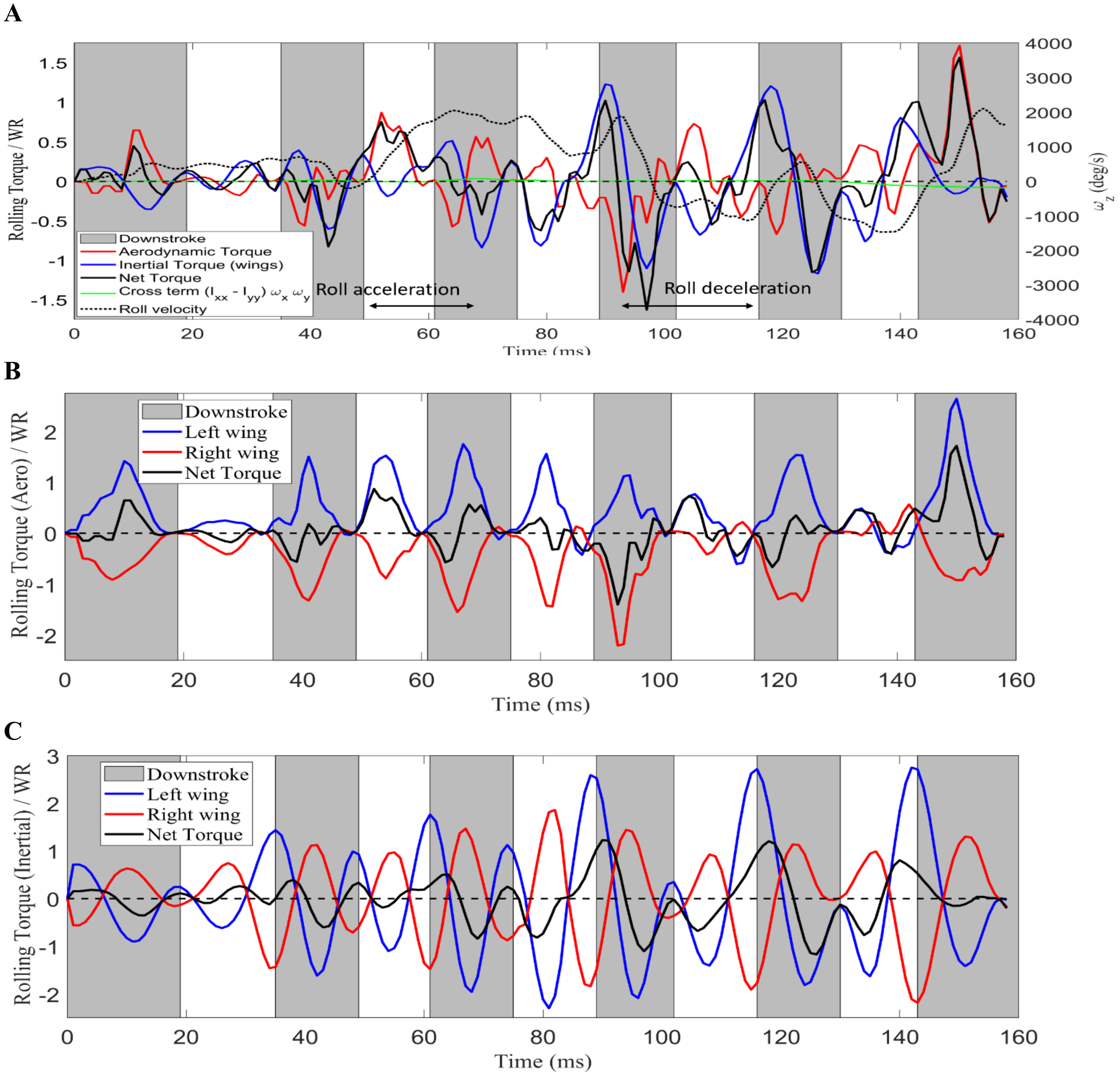


**Rolling torques (normalized by** ***WR*****) for Bird 2**. (**A**) Rolling torques generated by both wings. (**B**) Aerodynamic rolling torques produced by left and right wings**,** (**C**) Inertial rolling torques produced by left and right wings.

**Fig. S10.**

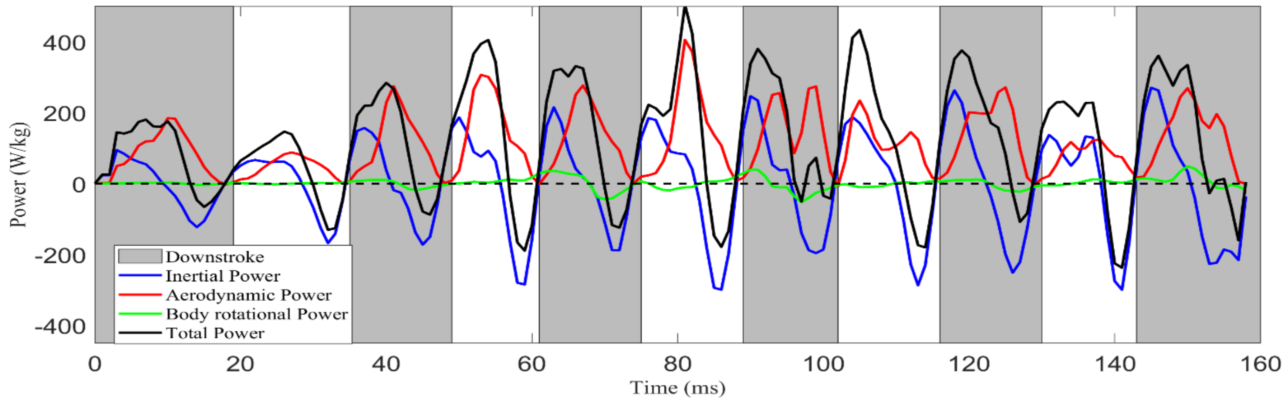


**Instantaneous mass-specific power for Bird 2, which includes the inertial power and aerodynamic power of the wings, and the body rotational power.**

**Fig. S11**

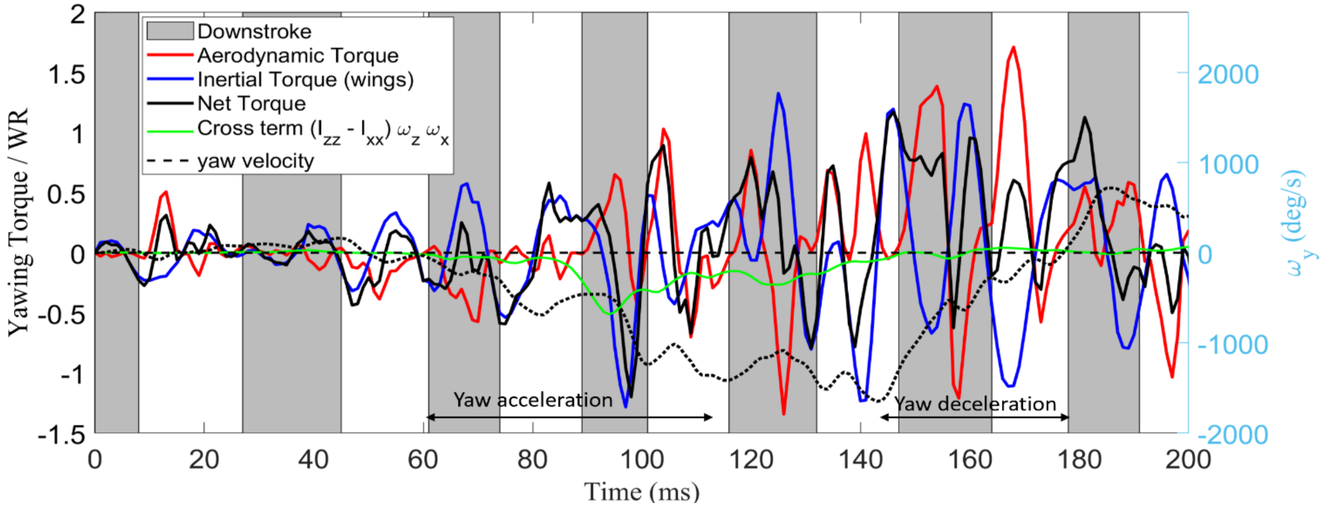


**Yawing torques (normalized by *WR*) for Bird 1.**

**Movie S1.**
Flow field from CFD simulation of Bird 1, where the vortex structures were visualized.

**Movie S2.**
Instant aerodynamic force vector with respect to the body for Bird 1, where the colors on the body surfaces represented the pressure distribution.

**Movie S3.**
Instant aerodynamic and inertial force vectors of the two wings from different view angles (vectors were plotted at wingtips for better visibility).
